# Prioritized imputed sequence variants from multi-population GWAS improve prediction accuracy for sea lice count in Atlantic salmon (*Salmo salar*)

**DOI:** 10.1101/2023.07.07.548148

**Authors:** Baltasar F. Garcia, Pablo A. Cáceres, Rodrigo Marín-Nahuelpi, Paulina Lopez, Daniela Cichero, Jorgen Ødegård, Thomas Moen, José M. Yáñez

## Abstract

Sea lice infestation is one of the major fish health problems during the grow-out phase in Atlantic salmon (*Salmo salar*) aquaculture. In this study, we integrated different genomic approaches, including whole-genome sequencing (WGS), genotype imputation and meta-analysis of genome-wide association studies (GWAS), to identify single-nucleotide polymorphisms (SNPs) associated with sea lice count in Atlantic salmon. Different sets of trait-associated SNPs were prioritized and compared against randomly chosen markers, based on the accuracy of genomic predictions for the trait. Lice count phenotypes and dense genotypes of five breeding populations challenged against sea lice were used. Genotype imputation was applied to increase SNP density of challenged animals to WGS level. The summary statistics from GWAS of each population were then combined in a meta-analysis to increase the sample size and improve the statistical power of associations. Eight different genotyping scenarios were considered for genomic prediction: 70K_array: 70K standard genotyping panel; 70K_priori: 70K SNPs with the highest p-values identified in the meta-analysis; 30K_priori: 30K SNPs with the highest p-values identified in the meta-analysis; WGS: SNPs imputed to whole-genome sequencing level; and the remaining four scenarios were the same SNP sets with a linkage disequilibrium (LD) pruning filter: 70K_array_LD; 70K_priori_LD; 30K_priori_LD and WGS_LD, respectively. Genomic prediction accuracy was evaluated using a five-fold cross-validation scheme in two different populations excluding them from the meta-analysis to remove possible validation-reference bias. Results showed significant genetic variation for sea lice counting in Atlantic salmon across populations, with heritabilities ranging from 0.06 to 0.24. The meta-analysis identified several SNPs associated with sea lice resistance, mainly in *Ssa03* and *Ssa09* chromosomes. Genomic prediction using the GWAS-based prioritized SNPs showed higher accuracy compared to using the standard SNP array in most of scenarios, achieving up to 57% increase in accuracy. Accuracy of prioritized scenarios was higher for the 70K_priori in comparison to 30K_priori. The use of WGS data in genomic prediction presented marginal or negative accuracy gain compared to the standard SNP array. The LD-pruning filter presented no benefits, reducing accuracy in most of scenarios. Overall, our study demonstrated the potential of prioritized of imputed sequence variants from multi-population GWAS meta-analysis to improve prediction accuracy for sea lice count in Atlantic salmon. The findings suggest that incorporating WGS data and prioritized SNPs from GWAS meta-analysis can accelerate the genetic progress of selection for polygenic traits in salmon aquaculture.

## 1. Introduction

One of the main sanitary problems in the Atlantic salmon (*Salmo salar)* production during marine grow-out phase is the infestation caused by sea lice. This ectoparasite causes huge economic losses in the salmon industry due to reduction of growth rate, increase of production costs associated to treatments and prevention of infestations, depreciation of filet quality and may represent a gateway for secondary infections increasing mortality rates (Brooker et al., 2018; Costello, 2009). In Chile, *C. rogercresseyi* is the main sea lice parasite affecting farmed Atlantic salmon, and costs associated to treatments may add around US$1.4 per fish kilogram on the final production cost (Dresdner et al., 2019).

An approach to reduce sea lice load is the implementation of breeding programs that select for genetically resistant fish to this parasite. Some studies showed significant genetic variation for *C. rogercresseyi* counting in Atlantic salmon with heritabilities (h²) ranging from 0.1 to 0.34 (Lhorente et al., 2012; Yáñez et al., 2014). These results were obtained using controlled infestation challenges to record lice count (LC) and pedigree information showing that a decrease of sea lice burden may respond to genetic selection in Atlantic salmon. Currently, aquaculture breeding programs have also included genomic information of animals to select candidates with higher accuracy (Yáñez et al., 2022). This has been possible due to advances in the discovery and characterization of single nucleotide polymorphisms (SNPs), by means of high-throughput sequencing and genotyping technologies. This information can be applied to perform genome-wide association studies (GWAS) allowing the identification of genetic markers significantly associated with the target traits (Goddard and Hayes, 2009). In addition, genomic selection (GS) may also be implemented using SNP information. GS consists of estimating genomic breeding values (GEBVs) of selection candidates, with or without phenotype information, using a related reference population with genotypes and phenotypes through a prediction equation (Meuwissen et al., 2001). GWAS have been performed for *C. rogercresseyi* counting in Atlantic salmon using a 50K SNP array revealing a polygenic architecture for this trait, i.e., several genes of small effects affecting the trait (Correa et al., 2017b; Robledo et al., 2019). GS method has also been employed for the same trait and compared to predictions using only pedigree information (Correa et al., 2017a; Tsai et al., 2017, 2016). These approaches showed higher accuracy of prediction when genomic information is included, achieving a 20-30% increase in comparison to pedigree-based genetic evaluations.

Several factors may affect the accuracy of genomic prediction, e.g., heritability and genetic architecture of trait, number of genotyped animals, SNP density, among others (Goddard et al., 2010; Zhang et al., 2019). In this regard, it is theoretically expected that the use millions of SNPs from whole-genome sequencing (WGS) increases genomic prediction accuracy because it potentially may include the causal mutations affecting the trait, in comparison to SNP arrays that use thousands of pre-defined SNPs (MacLeod et al., 2016). However, in practical terms the use of WGS in genomic prediction has two main drawbacks: higher cost and a relatively small accuracy gain in comparison to SNP arrays (Pérez-Enciso et al., 2015). The cost issue can be partially solved by using genotype imputation. Genotype imputation is performed using a small group of sequenced animals as a reference to predict the missing genotypes of target animals genotyped using lower density of SNPs (Calus et al., 2014; Huang et al., 2012). The marginal gain in accuracy might be related to the introduction of “noise” into predictions, due to the high number of SNPs that do not affect the target trait causing overparameterization (Raymond et al., 2018; van den Berg et al., 2016). This effect may be reduced if WGS data is filtered, by prioritizing only potential SNPs truly affecting the traits. GWAS results using WGS data, for example, could help in this task by selecting the most relevant trait-associated SNPs and using them on genomic prediction.

The objectives of this study were to: i) perform a meta-analysis of GWAS for lice count using thousands of animals from different populations and imputed WGS genetic variants, ii) estimate the accuracy of genomic prediction using sets of prioritized SNPs based on meta-analysis GWAS results, and iii) compare a regular predefined SNP array to different sets of prioritized SNPs.

## 2. Material and methods

### 2.1 Animals and sea lice phenotypes

Animals used in this study belong to the AquaGen breeding company. A total of five year-classes (herein called Pop1, Pop2, Pop3, Pop4 and Pop5) were challenged against *C. rogercresseyi*. All challenges were performed similarly with slight changes regarding day-post-infestation and frequency of sea lice counting. Briefly, animals identified with PIT-tags were received in a tank of 7 m^3^ at 10 ppt of salinity and 12°C during 12 days for acclimatization with gradual increase of salinity until reaching 100% seawater. Then, fish were challenged with *C. rogercresseyi* copepodites obtained from parasite ovigerous females. A total of 35 copepodites were infested per fish. After an infestation period, lice count (LC) phenotype, weight at counting and PIT-tag were recorded for each animal. After first infestation and measurement of LC, animals were deparasitized by gradually reducing the salinity until 5 ppm, and then, salinity was re-stablished to the previous level. A second infestation and LC measurement was performed using the same protocol described previously. Population 1, 2, 3 and 4 had two LC measurement timepoints. Pop 4 also was challenged in two different tanks (tk1 and tk2). Finally, Pop5 had three LC measurement timepoints (t1, t2 and t3) in each one of the two challenges, summing up six LC phenotypes. Table 1 summarizes the number of challenged animals, days at LC and descriptive statistics of LC and weight for each population.

**Table 1.** Summary of descriptive statistics for day counting, weight and lice count of Atlantic salmon populations challenged for sealice susceptibility.

| Pop1 (N<sup>a</sup>= 1,369) |  |  |  |  |  |  |
| --- | --- | --- | --- | --- | --- | --- |
| Trait<sup>b</sup> | Day<sup>c</sup> | Mean weight (sd)<sup>d</sup> | Mean LC (sd)<sup>e</sup> | Min<sup>f</sup> | Max<sup>g</sup> | CV (%)<sup>h</sup> |
| LC1 | 17 | 187.6 ± 47.6 | 30 ± 11.5 | 6 | 89 | 38.3 |
| LC2 | 18 | 307 ± 87.7 | 26.1 ± 9.8 | 4 | 70 | 37.6 |
| Pop2 (N= 1,926) |  |  |  |  |  |  |
| LC1 | 10 | 195.2 ± 50.8 | 26.3 ± 9.1 | 4 | 79 | 34.6 |
| LC2 | 8 | 214.1 ± 61.2 | 27.7 ± 8.8 | 5 | 66 | 31.8 |
| Pop3 (N= 1,512) |  |  |  |  |  |  |
| LC1 | 13 | 123.8 ± 29.6 | 31 ± 15.7 | 3 | 132 | 50.5 |
| LC2 | 13 | 214.1 ± 46.3 | 27.2 ± 15.1 | 0 | 134 | 55.6 |
| Pop4 (N= 1,310) |  |  |  |  |  |  |
| LC1_tk1 | 2 | 271.2 ± 65.6 | 10 ± 6 | 0 | 60 | 60.5 |
| LC1_tk2 | 3 | 288.2 ± 75.6 | 17.1 ± 9 | 1 | 74 | 52.5 |
| LC2_tk1 | 12 | 440.3 ± 120.6 | 18.6 ± 7.7 | 3 | 68 | 41.4 |
| LC2_tk2 | 12 | 485.6 ± 134.2 | 32 ± 11.8 | 5 | 90 | 37.1 |
| Pop5 (N=1,127) |  |  |  |  |  |  |
| LC1_t1 | 4 | 169.1 ± 24.1 | 18 ± 8.2 | 2 | 75 | 45.8 |
| LC1_t2 | 8 | 166.9 ± 23.8 | 16.4 ± 8.1 | 4 | 107 | 49.1 |
| LC1_t3 | 15 | 170.6 ± 28.5 | 18.9 ± 8.7 | 2 | 85 | 45.7 |
| LC2_t1 | 4 | 179 ± 29.9 | 19.6 ± 7.5 | 4 | 99 | 38.2 |
| LC2_t2 | 10 | 188.1 ± 32.8 | 17 ± 7.7 | 3 | 123 | 45.5 |
| LC2_t3 | 15 | 184 ± 31.7 | 17.4 ± 6.7 | 4 | 57 | 38.8 |

<sup>a</sup>Number of animals
<sup>b</sup>Trait: lice count at different time points. LC1, LC2 represents first and second infection, respectively. tk1 and tk2 represent lice counting in two different tanks, respectively. t1, t2 and t3 represent the sequential counting at different time points within same challenge, respectively.
<sup>c</sup>Day of counting after infestation
<sup>d</sup>Mean weight of challenged animals (standard deviation)
<sup>e</sup>Mean lice count (standard deviation)
<sup>f</sup>Minimum lice count
<sup>g</sup>Maximum lice count
<sup>h</sup>Coefficient of variation for lice count

### 2.2 Genotyping

Animals of all challenged populations (N= 7,244) were genotyped using low-density SNP arrays varying from 50, 60 and 70K SNPs. These were AquaGen’s customized SNP arrays applied for routine genomic evaluations and updated periodically by including relevant SNPs associated to economically important traits. Additionally, genotype data from other three different populations were used as a reference for genotype imputation. These populations were comprised of animals from previous generations that originated the recent populations challenged for sea lice. These three reference populations were genotyped using a 200K ThermoFisher (Affymetrix) SNP array (Ref_MD), 930K ThermoFisher (Affymetrix) SNP array (Ref_HD), and whole-genome sequencing (Ref_WGS). Quality control (QC) of genotypes was performed independently for all five challenged and three references populations according to the following filters: call-rate for SNPs < 0.9, minor allele frequency (MAF) < 0.01 and Hardy-Weinberg equilibrium (HWE) (Bonferroni corrected p-value < 0.05). Number of animals and SNPs of reference populations are available in Table 2.

**Table 2.** Number of animals and SNPs for all Atlantic salmon populations challenged against sealice and used as reference in the genotype imputation to WGS level.

| Pop.<sup>1</sup> | N_Ind.<sup>2</sup> | N_SNPs<sup>3</sup> | Imputation (MD)<sup>4</sup> | Imputation (HD)<sup>5</sup> | Imputation (WGS)<sup>6</sup> | Post-QC<sup>7</sup> |
| --- | --- | --- | --- | --- | --- | --- |
| Pop1 | 1,369 | 47,374 | 208,347 | 600,196 | 2,900,645 | 2,509,677 |
| Pop2 | 1,926 | 46,214 | 208,347 | 600,196 | 2,865,066 | 2,301,569 |
| Pop3 | 1,512 | 53,512 | 209,060 | 600,491 | 2,865,580 | 2,307,797 |
| Pop4 | 1,310 | 59,213 | 227,179 | 616,898 | 2,907,072 | 2,436,326 |
| Pop5 | 1,127 | 61,781 | 221,882 | 613,138 | 2,969,396 | 2,507,142 |
| Ref_MD | 1,480 | 215,379 | - | - | - | - |
| Ref_HD | 1,314 | 633,316 | - | - | - | - |
| Ref_WGS | 299 | 4,277,345 | - | - | - | - |

<sup>1</sup>Populations. Pop1 to Pop5 are the populations challenged against sealice and genotyped at low density (50, 60 and 70k). Ref\_MD, Ref\_HD and Ref\_WGS are animals from previous generations genotyped at medium (200K), high (930K) and whole-genome sequencing level, respectively, used as reference in the step-wise genotype imputation
<sup>2</sup>Number of animals
<sup>3</sup>Number of SNPs after quality control of genotypes prior to imputation. call-rate < 0.9, minor allele frequency (MAF) < 0.01 and Hardy-Weinberg equilibrium (p-value < 0.05/number of SNPs remaining after the two previous filters)
<sup>4</sup>Number of SNPs after imputation to 200k using only SNPs with accuracy of imputation greater than 0.8
<sup>5</sup>Number of SNPs after imputation to 930k using only SNPs with accuracy of imputation greater than 0.8
<sup>6</sup>Number of SNPs after imputation to WGS using only SNPs with accuracy of imputation greater than 0.8
<sup>7</sup>Number of SNPs after quality control of imputed genotypes. The same filter parameters used before were applied in this step, except for MAF that was changed to 0.05

### 2.3 Genotype imputation

A sequential “step-wise” imputation strategy was adopted to impute all challenged populations to WGS level, i.e., from low-density (Pop1, 2, 3, 4 and 5 genotyped with 50k to 70K) to 200K (Ref_MD), then from Ref_MD to 930K (Ref_HD) and finally from Ref_HD to sequence level (Ref_WGS). Validations of genotype imputation were performed for all reference populations to remove SNPs with low accuracy of imputation. This was done by masking 20% of reference animals in five different cross-validation groups. The masked animals had only part of SNPs available, and these SNPs were in common to all challenged populations. For example, in the validation of 200K imputation, a set of 43,761 SNPs common SNPs segregating among all five challenged populations (Pop 1 to Pop5) was used as target SNPs. Imputation was performed using FImpute3 software (Sargolzaei et al., 2014) using the remaining 80% animals as reference in each group. Imputation accuracy was assessed by Pearson’s coefficient of correlation between imputed and observed (originals) genotypes at SNP and individual level. SNPs with imputation accuracy lower than 0.8 were removed from the final round of imputation of challenged populations. This process was repeated for Ref_HD and Ref_WGS populations until final imputation of all challenged populations to WGS level. After imputation, a new QC of genotypes was applied using the same filters described before, except for MAF that was changed to > 0.05 threshold. The final number of SNPs after imputation for all density levels is shown in Table 2.

### 2.4 GWAS for LC and meta-analysis

GWAS for LC at different timepoints were performed for each population separately using the final set of SNPs imputed to WGS level. The GCTA software (Yang et al., 2011) was used for GWAS and a genomic relationship matrix (GRM) was constructed to be included in the model to account for population relatedness. The model used was:

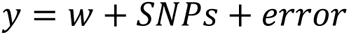

where *y* is LC, *w* is the weight at counting used as fixed effect, *SNPs* is the fixed SNP effect and *error* is the random residual. GWAS were performed using the GRM and the MLMA-LOCO function, which is based on an association analysis within the chromosome and the candidate SNP is located and excluded from calculating the GRM in each iteration step. The heritability was also estimated for each separate single-population GWAS analysis using REML procedures implemented in GCTA.

The summary statistics from GWAS carried out on each population were combined and used to perform a meta-analysis. We opted to use the meta-analysis to improve statistical power of associations and prioritize variants associated to LC across different populations. The meta-analysis was conducted across all five populations considering all LC as the same trait due to high genetic correlation among them (results not shown). The METAL software (Willer et al., 2010) was used to implement the meta-analysis combining the p-values and direction of SNP effects from the individual GWAS based on the sample size to compute an overall Z-score:

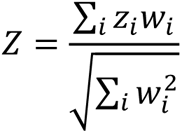

where *w*_*i*_ is the square root of the sample size of population *i*, and *z*_*i*_ is the Z-score for population *i* calculated as 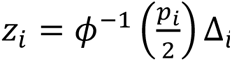, where *φ* is the cumulative distribution function, and *p*_*i*_ and Δ_*i*_ are the p-value and direction of effect for population *i*, respectively. The Z-score was then used to estimate an overall p-value weighting the effect size estimates by their estimated standard errors. The overall p-value was used to construct the sets of prioritized SNPs to be furtherly used in the genomic prediction analyses. A potential bias when using prioritized SNPs from GWAS into genomic prediction is to prioritize and validate genomic prediction in the same population (Yoshida and Yáñez, 2021). To avoid this effect, Pop4 and Pop5 were removed from meta-analysis when genomic prediction was performed in these populations, i.e., genomic prediction for Pop4 used only prioritized SNPs from meta-analysis using Pop1, 2, 3 and 5. The same approach was used when genomic prediction was performed for Pop5, which used only SNPs prioritized from meta-analysis using Pop1, 2, 3 and 4.

### 2.5 Prioritized SNP sets and genomic prediction

Genomic prediction was evaluated using eight different SNP scenarios (Table 3): 70K_array: 70K standard genotyping panel; 70K_priori: 70K SNPs with the highest p-values identified in the meta-analysis; 30K_priori: 30K SNPs with the highest p-values identified in the meta-analysis; WGS: SNPs imputed to whole-genome sequencing level. The other four scenarios included the same SNP sets with a linkage disequilibrium (LD) pruning filter with a threshold r² < 0.2: 70K_array_LD: LD-pruned 70K standard genotyping panel; 70K_priori_LD: LD-pruned 70K SNPs with the highest p-values identified in the meta-analysis; 30K_priori_LD: LD-pruned 30K SNPs with the highest p-values identified in the meta-analysis; WGS_LD: LD-pruned SNPs imputed to whole-genome sequencing level. LC at different days were used as phenotypes in both prediction populations (four different counting days for Pop4 and six different counting days for Pop5).

**Table 3.** Number of SNPs for each genomic prediction scenario evaluated in populations Pop4 and Pop5 challenged against sealice.

| Scenario<sup>1</sup> | Pop4<sup>2</sup> | Pop5<sup>3</sup> |
| --- | --- | --- |
| 70K_array | 59,213 | 61,781 |
| 70K_array_LD | 44,662 | 41,572 |
| 70K_priori | 70,000 | 70,000 |
| 70K_priori_LD | 26,458 | 24,317 |
| 30K_priori | 30,000 | 30,000 |
| 30K_priori_LD | 8,990 | 14,953 |
| WGS | 2,675,940 | 2,507,142 |
| WGS_LD | 85,810 | 162,331 |

<sup>1</sup>Scenarios: 70K\_array: 70K standard genotyping panel; 70K\_array\_LD: LD-pruned 70K standard genotyping panel ; 70K\_priori: 70K SNPs with the highest p-values identified in the meta-analysis; 70K\_priori\_LD: LD-pruned 70K SNPs with the highest p-values identified in the meta-analysis; 30K\_priori: 30K SNPs with the highest p-values identified in the meta-analysis; 30K\_priori\_LD: LD-pruned 30K SNPs with the highest p-values identified in the meta-analysis; WGS: SNPs imputed to whole-genome sequencing level; WGS\_LD: LD-pruned SNPs imputed to whole-genome sequencing level
<sup>2</sup>Pop4: Population used to evaluate genomic prediction accuracy with two lice count phenotypes in two different tanks
<sup>3</sup>Pop5: Population used to evaluate genomic prediction accuracy with three lice count phenotypes in two sequential challenges

The genomic prediction accuracy was evaluated in Pop4 and Pop5 using a five-folded cross-validation scheme in which 80% of animals were used as training set to estimate SNP effects and the remaining 20% were used as validation. The genomic best linear unbiased predictor (GBLUP) method was used to estimate the genomic breeding values (GEBVs). The following model was used in the BLUPF90 software (Misztal et al., 2018):

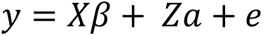

where *y* is a vector of phenotypes (LC at different days), β is a vector of fixed effects (body weight at LC measurement), *a* is a vector of random additive polygenic genetic effects that follows a normal distribution 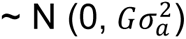, where *G* is the genomic relationship matrix (VanRaden, 2008) and 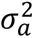 is the additive genetic variance *X* and *Z* are incidence matrices, *e* is the random residual error with a distribution 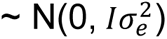, where *I* is an identity matrix and 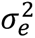 is the residual variance. The accuracy of genomic prediction was assessed by dividing the Pearson’s correlation coefficient between GEBVs and the phenotypes by the square root of heritability estimated using the standard 70K SNP panel.

## 3. Results

### 3.1 Phenotypes, genotypes and imputation

A large phenotypic variation was observed across all five challenges with coefficient of variation for LC ranging from 31.8 in Pop2 to 60.5 in Pop4. The differences in LC coefficient of variation most likely happened due to different days of LC in each challenge ranging from 2 to 18 days after infestation (Table 1).

After genotype QC, the 7,244 challenged animals that were genotyped using low-density SNP arrays passed the filters, showing from 46,214 available SNPs in Pop2 to 61,781 SNPs in Pop5 (Table 2). These differences occurred due to different SNP array densities (i.e., from 50 to 70K) used in different populations, as they were updated periodically to include additional SNPs for genomic prediction purposes. Overall, validation of genotype imputation yielded high accuracy with 98.1, 91 and 57% of SNPs imputed with accuracy above 0.8, for medium-density (200K), high-density (930K) and WGS level, respectively (Supplementary 1). After QC, all challenged populations had around 2.5M SNPs imputed to WGS level with high accuracy.

### 3.2 GWAS and meta-analysis

Heritability explained by SNPs imputed to WGS level is presented in Table 4. All LC phenotypes presented significative and low to moderate heritability values ranging from 0.06 to 0.24. These results reinforce the idea that lice burden presents sufficient genetic variance and could respond to selection in Atlantic salmon.

**Table 4.**
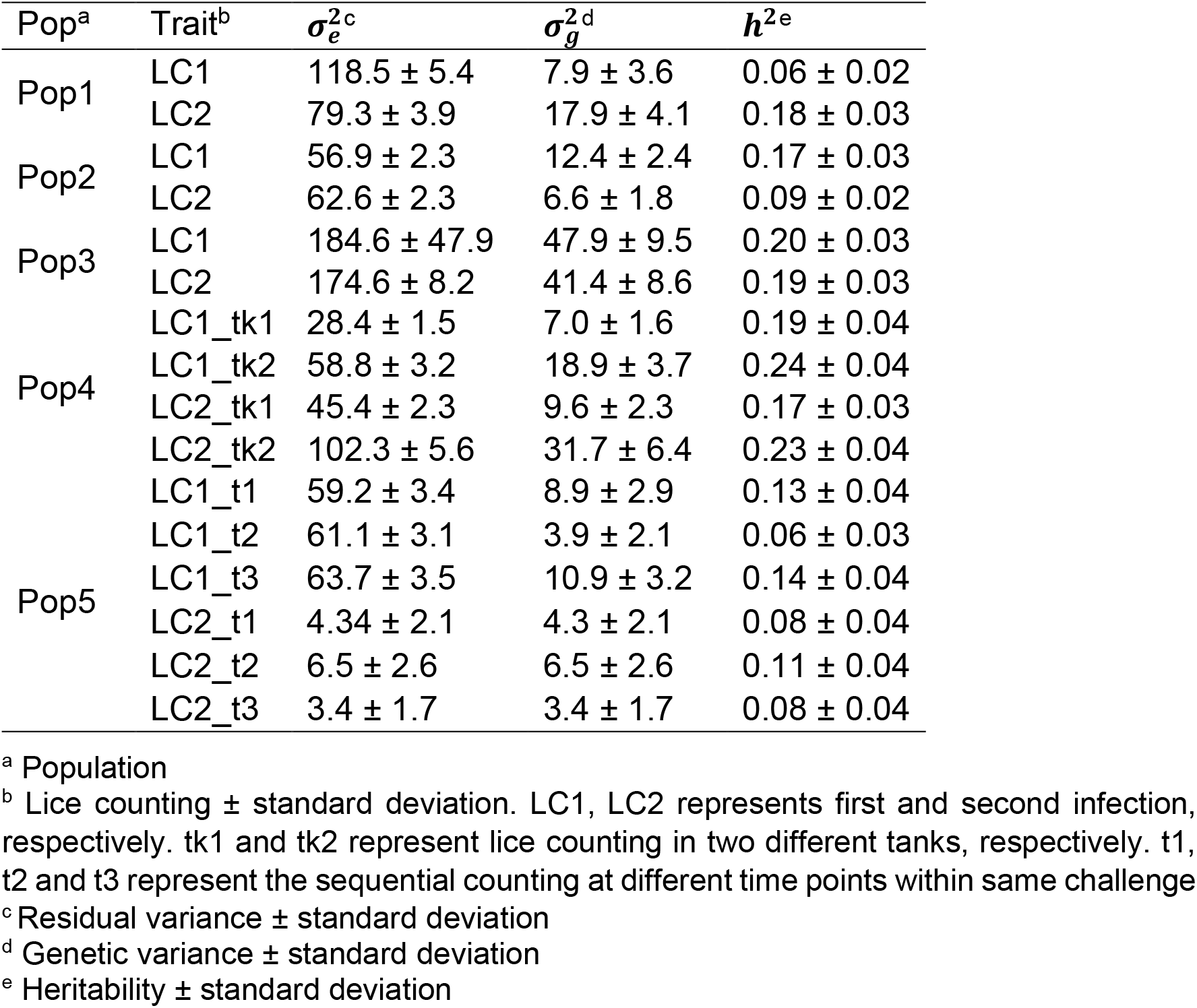
Genetic parameters for all Atlantic salmon populations challenged against sealice evaluated in the meta-analysis and genomic prediction study using WGS imputed data.

Single-trait GWAS results did not show any significant peaks in any population evaluated (Supplementary 2). However, when the meta-analysis was performed, the overall p-value increased substantially, and we were able to identify significant associations in *Ssa3* and *Ssa9* chromosomes (Figure 1). This is the main benefit of meta-analysis approach: to increase power to detect associations by overlapping and scaling different GWAS results. To avoid bias in accuracy estimation, genomic prediction using prioritized SNPs were performed based on meta-analysis results excluding the validation populations (i.e., Pop4 or Pop5). The Manhattan plots of the GWAS meta-analyses results are available in Supplementary 3.

**Figure 1.**
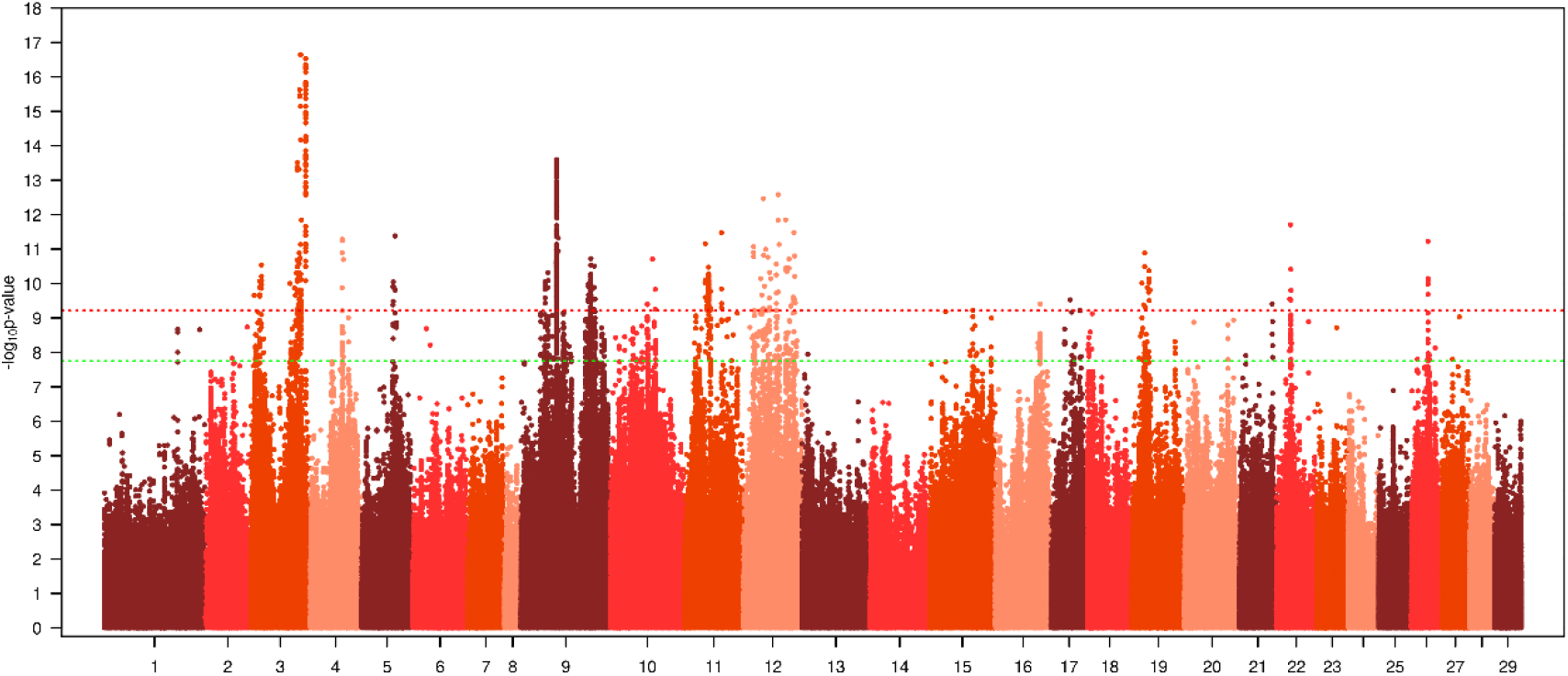
Manhattan plot of meta-analysis of GWAS results for lice count in Atlantic salmon using genotypes imputed to WGS level from all five populations (Green and red lines indicate chromosome - and genome-wide significance thresholds, respectively).

### 3.3 Genomic prediction accuracy

Genomic prediction using prioritized SNPs presented higher accuracy in most of the LC timepoints for both validation populations. For Pop4 (Table 5), when using the 70K SNPs with highest p-values from meta-analysis and LD pruning of markers, the increase in accuracy was observed in six out of eight LC timepoints ranging from 0.69 to 55.81% considering all LC timepoints and both challenge tanks. In only two LC timepoints, LC2_tk1 and LC2_tk2, there was a slight decrease in the accuracy of genomic prediction, -0.65 and -5.73%, respectively, for the 70K_priori scenario. For the 30K_priori and 30K_priori_LD, the prioritization was less effective (half of LC timepoints showed increase in accuracy) with higher increase ranging from 6.15 to 57.82% and decrease from -0.35 to -9.97% in comparison to 70K_array scenario. The scenarios with highest number of SNPs (WGS and WGS_LD) did not reflect in higher accuracy and only for LC1_tk2 there was a substantial increase in accuracy (37.67 and 23.31%, for WGS and WGS_LD respectively). A similar trend was observed in Pop5 (Table 6) regarding the same scenarios evaluated: 70K_priori and 70K_priori_LD presented increase in accuracy in eleven out of twelve LC timepoints evaluated in this population (from 1.62 to 25.17%), 30K_priori and 30K_priori_LD were less effective, and WGS scenarios were the worst scenarios in terms of accuracy.

**Table 5.** Accuracy of genomic prediction for lice count (LC) in Pop4 using different SNP scenarios.

| Trait<sup>1</sup>/Scenario<sup>2</sup> | LC1_tk1 | LC1_tk2 | LC2_tk1 | LC2_tk2 |
| --- | --- | --- | --- | --- |
| 70K_array | **0.51±0.07 (0%)** | **0.36±0.07 (0%)** | **0.43±0.07 (0%)** | **0.70±0.26 (0%)** |
| 70K_array_LD | **0.28±0.17 (-44.66%)** | **0.31±0.20 (-16.37%)** | **0.43±0.06 (1.29%)** | **0.70±0.27 (0.70%)** |
| 70K_priori | **0.52±0.09 (1.72%)** | **0.57±0.13 (55.81%)** | **0.43±0.07 (-0.65%)** | **0.66±0.29 (-5.73%)** |
| 70K_priori_LD | **0.53±0.07 (5.18%)** | **0.56±0.11 (53.08%)** | **0.45±0.05 (5.82%)** | **0.70±0.23 (0.69%)** |
| 30K_priori | **0.50±0.09 (-2.39%)** | **0.57±0.13 (57.07%)** | **0.43±0.06 (-0.35%)** | **0.63±0.28 (-9.97%)** |
| 30K_priori_LD | **0.54±0.07 (5.52%)** | **0.58±0.11 (57.82%)** | **0.46±0.04 (6.15%)** | **0.69±0.22 (-0.85%)** |
| WGS | **0.50±0.07 (-1.01%)** | **0.50±0.11 (37.67%)** | **0.42±0.06 (-2.21%)** | **0.70±0.27 (0.01%)** |
| WGS_LD | **0.40±0.06 (-14.1%)** | **0.45±0.07 (23.31%)** | **0.36±0.07 (-16.67%)** | **0.70±0.30 (1.22%)** |

<sup>1</sup>Traits: Lice count (LC) of two sequential challenges (LC1 and LC2) in two different challenge tanks (tk1 and tk2)
<sup>2</sup>Scenarios: 70K\_array, 70K standard genotyping panel; 70K\_priori: 70K SNPs with the highest p-values identified in the meta-analysis; 30K\_priori: 30K SNPs with the highest p-values identified in the meta-analysis; WGS: SNPs imputed to whole-genome sequencing level. \_LD represents linkage disequilibrium pruning
\*Color scale in comparison to 70K\_array: light green (less than 5% increase), dark green (more than 5% increase), light red (less than 5% decrease) and dark red (more than 5% decrease)

**Table 6.** Accuracy of genomic prediction for LC in Pop5 using different SNP scenarios.

| Trait<sup>1</sup>/Scenario<sup>2</sup> | LC1_t1 | LC1_t2 | LC1_t3 |
| --- | --- | --- | --- |
| 70K_array | **0.37±0.17 (0%)** | **0.37±0.17 (0%)** | **0.31±0.03 (0%)** |
| 70K_array_LD | **0.35±0.09 (-5.46%)** | **0.39±0.10 (4.76%)** | **0.34±0.03 (7.14%)** |
| 70K_priori | **0.45±0.15 (20.4%)** | **0.40±0.12 (8.58%)** | **0.32±0.07 (1.62%)** |
| 70K_priori_LD | **0.45±0.15 (20.4%)** | **0.36±0.14 (-3.6%)** | **0.35±0.04 (10%)** |
| 30K_priori | **0.32±0.13 (-14.5%)** | **0.42±0.15 (13.78%)** | **0.32±0.09 (2.8%)** |
| 30K_priori_LD | **0.36±0.08 (-4.41%)** | **0.26±0.06 (-28.94%)** | **0.34±0.05 (8.37%)** |
| WGS | **0.33±0.07 (-11.09%)** | **0.37±0.09 (-1.75%)** | **0.31±0.03 (-1.17%)** |
| WGS_LD | **0.28±0.07 (-23.79%)** | **0.30±0.11 (-19.82%)** | **0.27±0.04 (-14.63%)** |
|  | LC2_t1 | LC2_t2 | LC2_t3 |
| 70K_array | **0.39±0.28 (0%)** | **0.30±0.12 (0%)** | **0.28±0.06 (0%)** |
| 70K_array_LD | **0.32±0.16 (-19.16%)** | **0.35±0.08 (13.67%)** | **0.27±0.05 (-3.6%)** |
| 70K_priori | **0.44±0.22 (12.57%)** | **0.34±0.07 (11.07%)** | **0.29±0.05 (4.8%)** |
| 70K_priori_LD | **0.46±0.24 (17.72%)** | **0.38±0.05 (25.17%)** | **0.31±0.07 (11.93%)** |
| 30K_priori | **0.35±0.15 (-11.47%)** | **0.34±0.07 (13.1%)** | **0.32±0.02 (12.66%)** |
| 30K_priori_LD | **0.39±0.08 (-1.14%)** | **0.38±0.05 (25.64%)** | **0.32±0.05 (13.56%)** |
| WGS | **0.31±0.13 (-21.04%)** | **0.33±0.08 (8.01%)** | **0.26±0.05 (-6.3%)** |
| WGS_LD | **0.29±0.02 (-26.21%)** | **0.30±0.04 (-1.3%)** | **0.24±0.07 (-14.5%)** |

<sup>1</sup>Traits: Lice count (LC) in two different challenges (LC1 and LC2) at three different days (t1, t2 and t3)
<sup>2</sup>Scenarios: 70K\_array, 70K standard genotyping panel; 70K\_priori: 70K SNPs with the highest p-values identified in the meta-analysis; 30K\_priori: 30K SNPs with the highest p-values identified in the meta-analysis; WGS: SNPs imputed to whole-genome sequencing level. \_LD represents linkage disequilibrium pruning
\*Color scale in comparison to 70K\_array: light green (less than 5% increase), dark green (more than 5% increase), light red (less than 5% decrease) and dark red (more than 5% decrease)

## 4. Discussion

In this study, we evaluated the effect of prioritization of SNPs on genomic prediction accuracy for lice count in Atlantic salmon. The prioritization was performed using the most relevant SNPs (lowest p-values for marker-trait association) revealed through GWAS performed based on WGS-imputed genotypes in multiple populations using a meta-analysis. For comparison purposes, we also evaluated the genomic prediction using a standard 70K genotyping array and WGS imputed data. Finally, LD-pruning was applied to all scenarios to decrease the number of SNPs and discard marker pairs that are high LD to avoid redundant genotypic information.

Phenotypic data from the five populations showed high variation. Despite differences regarding LC timepoints across populations, significative h² values (0.06 to 0.24) were estimated showing that there is a significant genetic component for lice burden in Atlantic salmon. Similar h² results were found for the same trait using pedigree and genomic data, with values ranging from 0.10 to 0.34 (Lhorente et al., 2012; Yáñez et al., 2014) and 0.11 to 0.12 (Correa et al., 2017a, 2016), for both sources of information, respectively. The differences between these estimations may be explained by: i) the different statistical methods implemented, as all studies reporting h² used BLUP approaches and we showed the SNP-heritability estimated using the GCTA software, ii) different levels of genetic variation for different populations, and iii) variation of environmental component among disease challenge experiments.

None SNP achieved statistical significance in any population when single-trait GWAS using WGS imputed was performed, becoming difficult to identify population-specific genomic regions associated to LC (Supplementary 2). Here, the implementation of meta-analysis represented an effective strategy to combine GWAS results from different populations to increase statistical power and estimate joint SNP effects across them (Begum et al., 2012). A potential drawback in the application of meta-analysis of GWAS is the possible use of different genotyping platforms across populations. Very often, there is low marker overlapping among SNP arrays, meaning that the number of common SNPs may be low. Here, this problem was solved using genotype imputation to WGS level, which in turn allowed to increase the SNP density to about 2.5M imputed markers. Despite genotype imputation errors may occur, we expected they happened at very low rates in the final imputed SNP set as shown by the validation results (Supplementary 1).

After meta-analysis, 936 SNPs were identified surpassing the chromosome-significance level (Figure 1). Two significant peaks were clearly identified at *Ssa03* and *Ssa09*. Cáceres et al. (2021) also found SNPs in the same chromosomes, both explaining low genetic variance for lice burden (around 3%) in a different population using a 50K SNP array. The *Ssa03* was also found associated to LC in the same population used by Cáceres et al. (2021) but using a different statistical method (Robledo et al., 2018). These results reinforce the idea that lice load has a polygenic architecture in Atlantic salmon, highlighting the importance of the implementation of GS to decrease lice burden via selective breeding.

With the recent advances in sequencing technologies and substantial reduction of costs of these approaches, it is expected that the use of WGS data is more commonly used to increase the accuracy of genomic prediction for breeding purposes. The main motivation is that WGS could potentially include the causal variants, maximizing prediction ability. However, most of the results obtained so far have been discouraging, showing similar or only marginal improvement in accuracy for several traits and species, such as reproduction and growth in dairy cattle (Frischknecht et al., 2018; Van Binsbergen et al., 2015), reproduction and production traits in pigs (Song et al., 2019), growth and carcass traits in chicken (Ye et al., 2019) and, growth traits under thermal stress in rainbow trout (Yoshida and Yáñez, 2021). This trend was also observed in our study with most of accuracies using WGS and WGS_LD being less effective than the standard 70K genotyping array. At least, three main reasons may be mentioned to explain this low WGS performance in genomic prediction models. First, due to genotype imputation, recombination events may be lost and the accuracy of imputation is usually low for rarer variants (i.e., large “gaps” close to a recombination region may not have sufficient information for the imputation) biasing estimation of genomic breeding values (Binsbergen et al., 2014; Garcia et al., 2022). Second, there is possibly a high number of non-causal variants in high linkage disequilibrium with causal mutations hardening the estimation of SNP effects explaining the genetic variance of traits (Heidaritabar et al., 2016). Third, the effectiveness of WGS in increasing genomic prediction accuracy is expected to be much higher when prior information on functional features is used in the set of pre-selected markers (Pérez-Enciso et al., 2015). Genotype imputation is difficult to improve unless a larger reference sample size is applied to impute rarer variants with higher accuracy (Druet et al., 2014). In this study, we tried to address the two last obstacles to maximize the incorporation of WGS into genomic prediction by pruning markers by LD and prioritizing of SNPs using meta-analysis of GWAS.

LD-pruning scenarios were performed to evaluate the decreasing of potential redundant SNPs on genomic prediction. LD is a primordial aspect of GS implementation as SNPs are expected to be in LD with at least on QTL affecting the trait and this effect across all genome may be able to capture the genetic variance (Meuwissen et al., 2001). The effect of LD-pruning reduced or slightly improved the accuracy of genomic prediction in comparison to non-LD-pruning scenarios when using low density of SNPs (70K_array, 70K_priori and 30K_priori). We expected that LD-pruning would have higher impact when using WGS information, but it was ineffective and, WGS_LD produced lower accuracies in comparison to WGS. Most likely, the remaining markers after LD-pruning presented imperfect LD with the causal mutations decreasing the prediction ability as showed by de los Campos et al. (2013). In addition, a possible confounding effect between LD and relationship among animals may have captured by the remaining SNPs hampering an increasing in the genomic prediction accuracy (Wientjes et al., 2013).

The prioritization of SNPs showed higher accuracy of genomic prediction in most of LC traits for both validation populations. The increase was more marked for LC1_tk2 in the Pop5 in which accuracy was of 53.08 to 57.82% higher compared with the four prioritized scenarios of SNPs. The reduction in the number of prioritized SNPs from 70K to 30K decreased accuracy in most scenarios. The increase of accuracy using prioritization of SNPs likely happened due to the meta-analysis of GWAS which provided valuable information about LC genetic architecture as more than 7,244 genotyped animals were included. Likely, the SNPs prioritized were the causal variants or were in high LD with the causal variants boosting the ability of prediction. Similar benefits were found in studies using the same prioritizing approach in dairy cattle (Raymond et al., 2018) and rainbow trout (Yoshida and Yáñez, 2021). However, other studies showed no benefit when using prioritization of SNPs from meta-analysis of GWAS results (Lu et al., 2020; Veerkamp et al., 2016). This inefficiency was also observed in some LC timepoints of the present study with an important accuracy variation across traits. For example, for LC1_tk2 in Pop4 and LC2_t2 in Pop5 practically all SNP scenarios presented higher accuracy in comparison to the standard 70K_array. In contrast, for LC2_tk2 in Pop4, for example, the increase was marginal for all scenarios or with slight reduction in accuracy. In spite there is high genetic correlations among traits, different lice counting timepoints may have caused this effect and the identification of the causal variants was variable across traits. This effect may be explained due to a random error caused by sampling as LC is subject to errors by the observer (counter) and size of sea lice (LC at early parasite stages of development is more difficult). It is worth mentioning that prediction accuracy was evaluated excluding the validation populations from meta-analysis providing more reliable estimations. A strategy to increase consistency of using prioritized SNPs on genomic prediction is to incorporate functional analysis as well, by integrating multi “omics” approaches to identify eQTLs, for example. The eQTLs involves studying the relationship between genetic variation and gene expression levels to identify regions in the genome that influence gene expression (Gilad et al., 2008). A “functional array” could increase the accuracy of genomic prediction as already demonstrated in *Drosophila melanogaster* (Ye et al., 2020) and sheep/dairy cattle (Daetwyler et al., 2019). Nevertheless, our study provides important results regarding the incorporation of WGS information into routine GS applications with substantial benefits in terms of accuracy when prioritized SNPs are applied.

## 5. Conclusion

We found a large phenotypic variation in LC across the five studied populations, with low to moderate heritability values, indicating a significant genetic component for sea lice burden in Atlantic salmon. Single-trait GWAS analysis did not yield any significant associations in individual populations. However, when a meta-analysis was performed, combining GWAS results from multiple populations, two significant associations were identified on chromosomes *Ssa03* and *Ssa09*. Genomic prediction using prioritized SNPs showed higher accuracy compared to standard genotyping arrays in most LC timepoints for the validation populations. Interestingly, the scenarios with the highest number of SNPs (whole-genome sequencing) did not necessarily result in higher accuracy. This suggests that the inclusion of all available SNPs, including potentially non-causal variants, may not necessarily improve prediction accuracy. Overall, these findings provide valuable insights into the application of genomic prediction for lice count in Atlantic salmon and contribute to the ongoing efforts to improve selective breeding strategies for sea lice resistance in this species.

## Acknowledgments

We acknowledge AquaGen Chile and Norway for providing the data and for their close collaboration. This research was performed under the collaborative project between the Aquaculture Genomics Laboratory of the University of Chile and AquaGen company.

## Author’s contribution

BFG: Conceptualization, Methodology, Formal analysis, Writing - Original Draft and Writing - Review & Editing. PAC: Formal analysis, Writing - Original Draft and Writing - Review & Editing. RMN: Formal analysis, Writing - Original Draft and Writing - Review & Editing. PL: Formal analysis and Writing - Review & Editing. DC: Writing - Review & Editing. JO: Writing - Review & Editing. TM: Writing - Review & Editing. JMY: Conceptualization, Methodology, Supervision and Writing - Review & Editing. All authors contributed to the design of the study, discussion of results and review of the manuscript. All authors read and approved the final version of the manuscript.

