## Supplemental 1 for "Prioritized imputed sequence variants from multi-population GWAS improve prediction accuracy for sea lice count in Atlantic salmon (*Salmo salar*)"

**Supplementary 1.** Accuracy of genotype imputation for 200K, 930K and WGS SNP densities at individual and SNP levels.

**Accuracy at Individual level**

| Correlation (r²) | Minimum | 1^st^ Quartile | Median | Mean | 3^rd^ Quartile | Maximum |
| --- | --- | --- | --- | --- | --- | --- |
| 200K | 0.94 | 0.98 | 0.98 | 0.98 | 0.99 | 1.00 |
| 930K | 0.36 | 0.96 | 0.97 | 0.96 | 0.97 | 0.99 |
| WGS | 0.55 | 0.65 | 0.67 | 0.67 | 0.69 | 0.80 |

**Accuracy at SNP level (Green line represents 0.8 imputation accuracy)**

**200K**

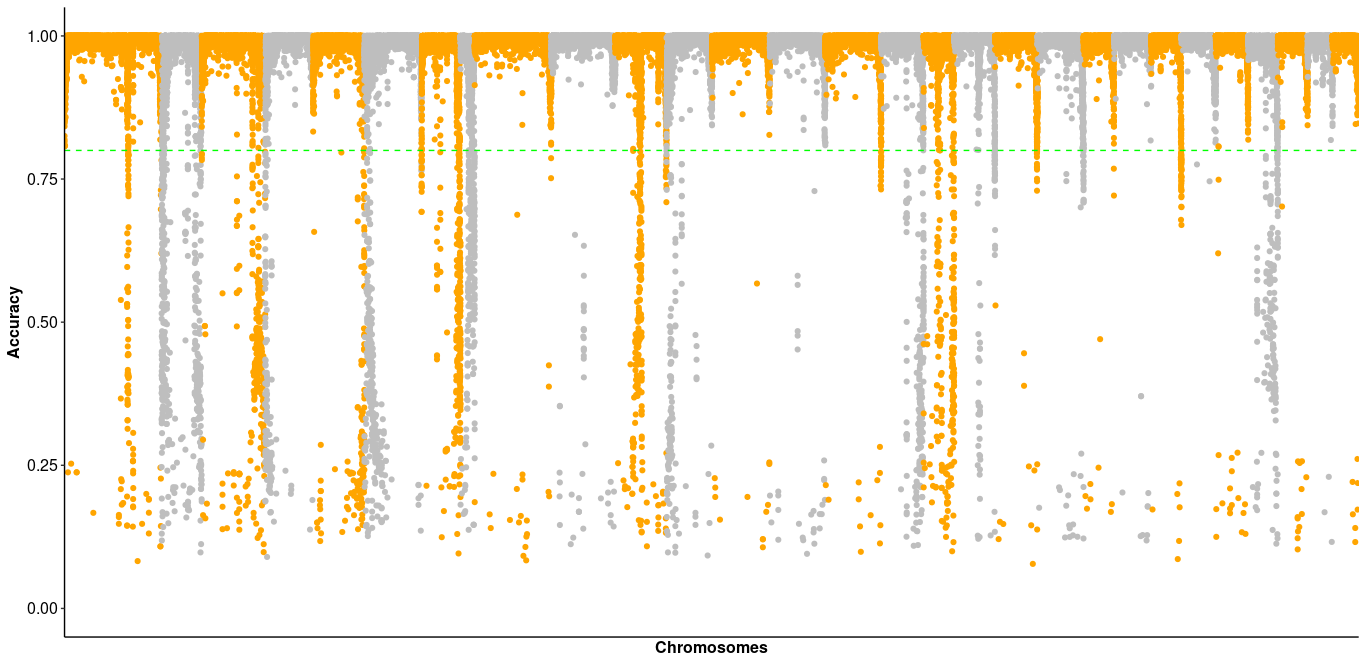

**930K**

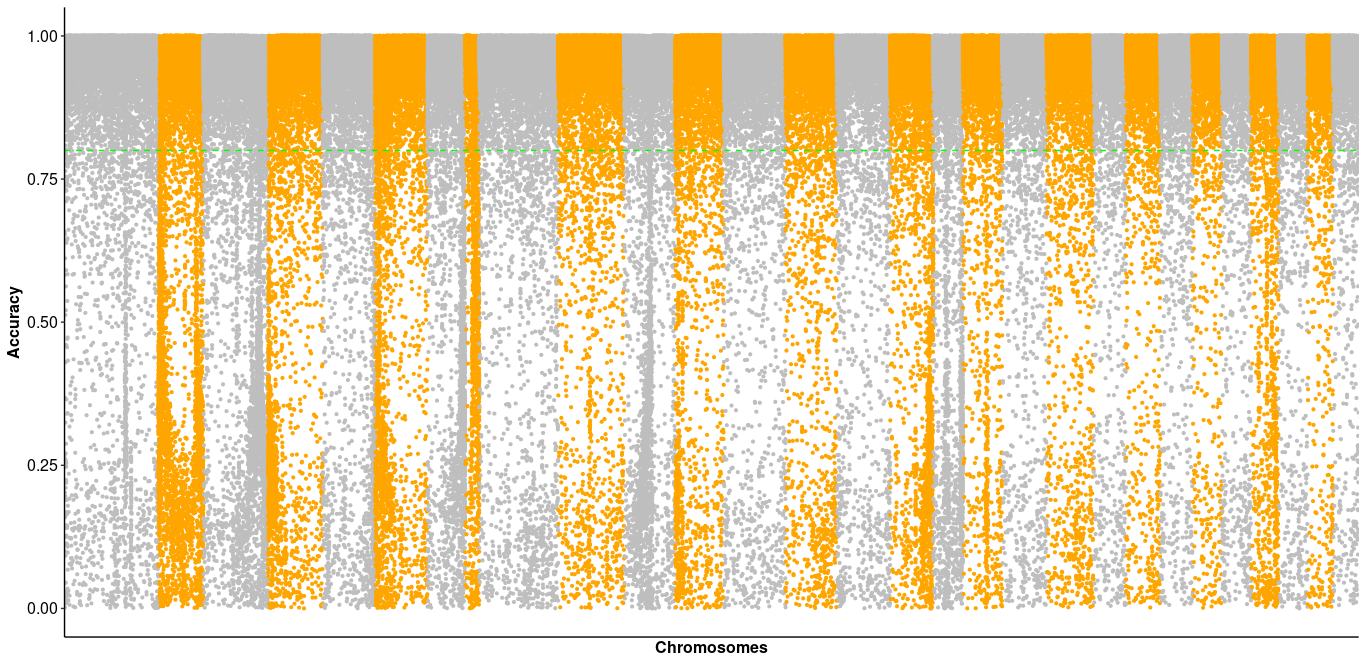

**WGS**

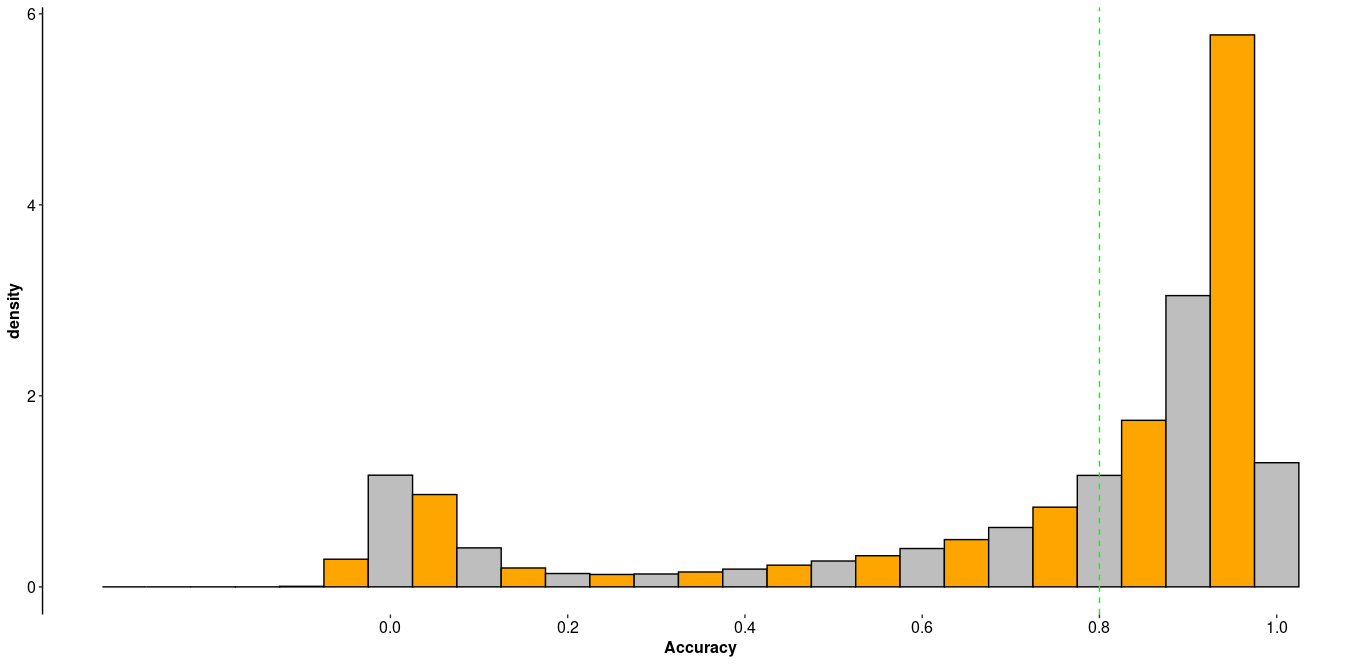
