## Supplemental 2 for "Prioritized imputed sequence variants from multi-population GWAS improve prediction accuracy for sea lice count in Atlantic salmon (*Salmo salar*)"

**Supplementary 2.** Individual GWAS for all populations and LC traits evaluated in this study.

**Pop1_LC1**

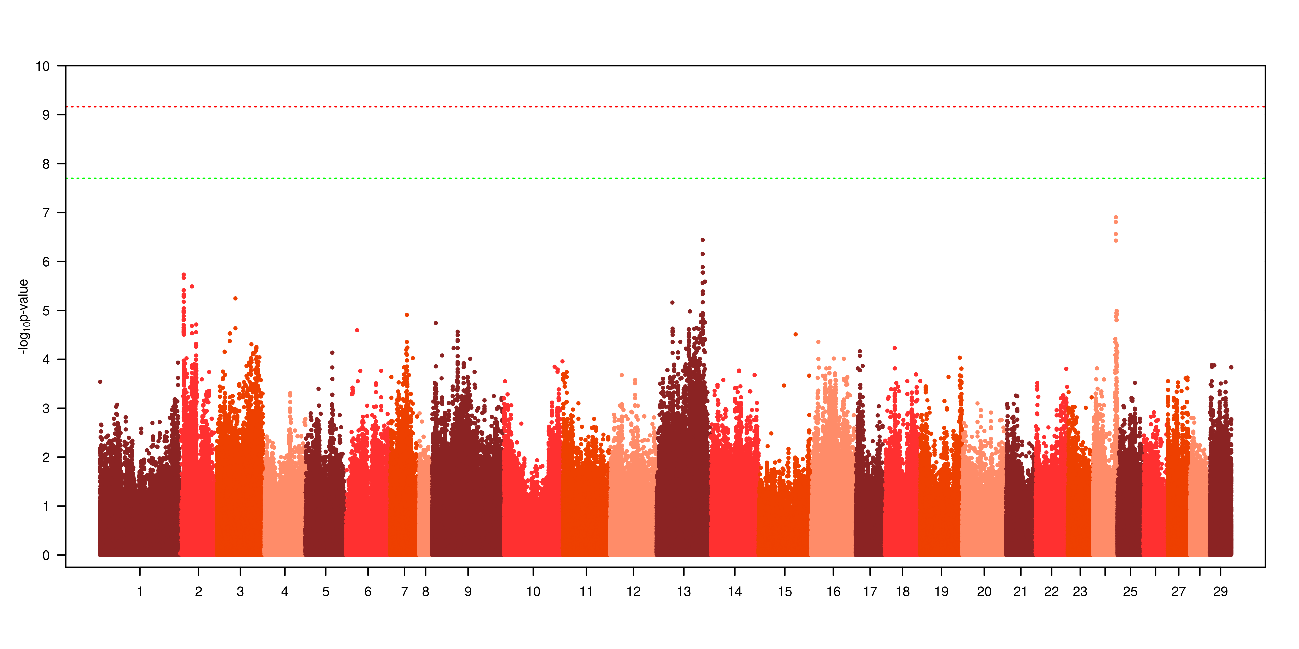

**Pop1_LC2**

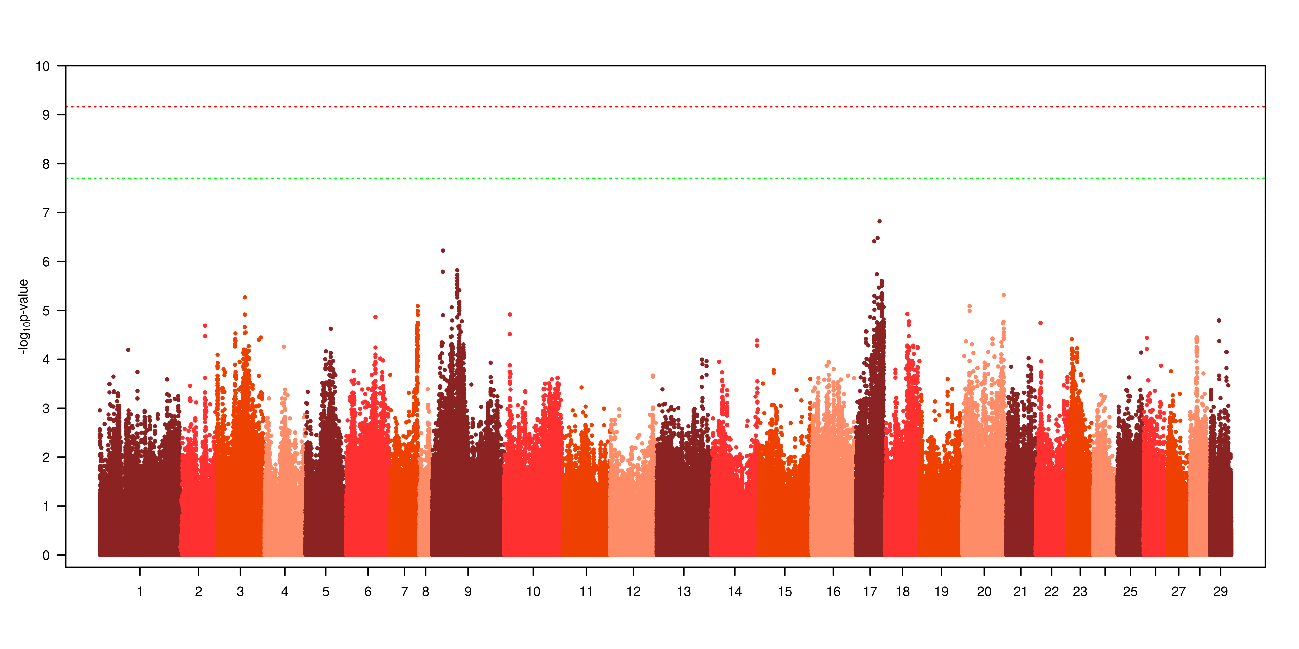

**Pop2_LC1**

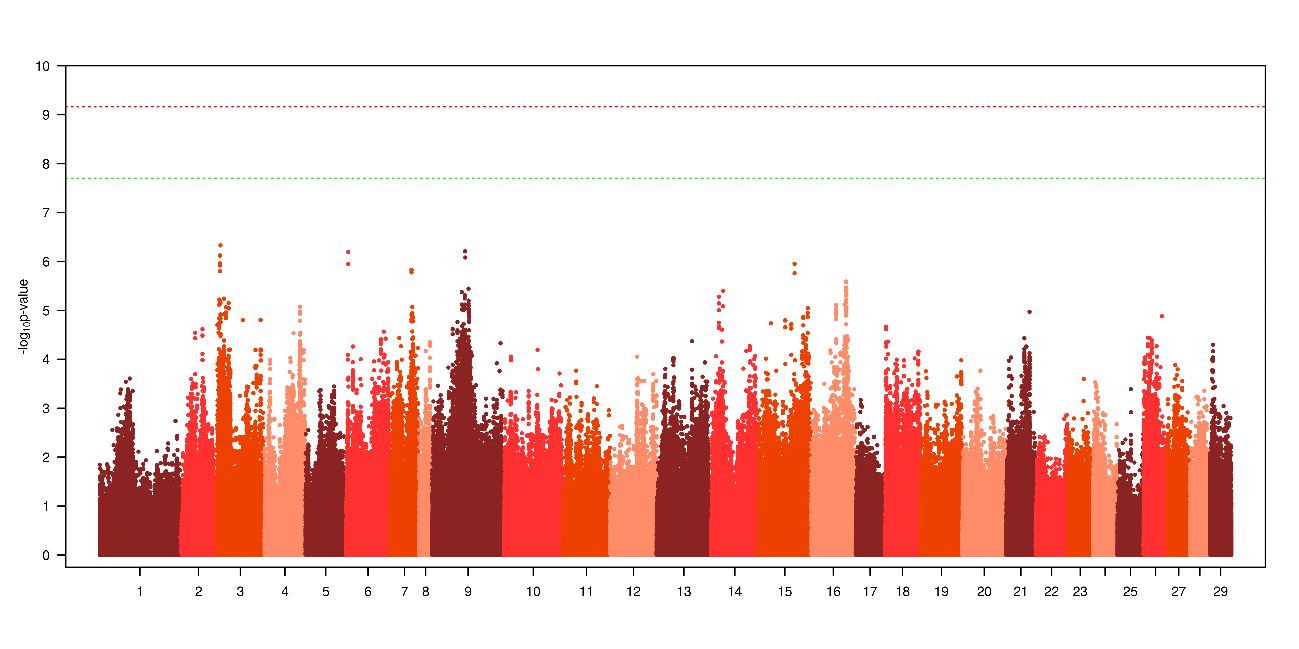

**Pop2_LC2**

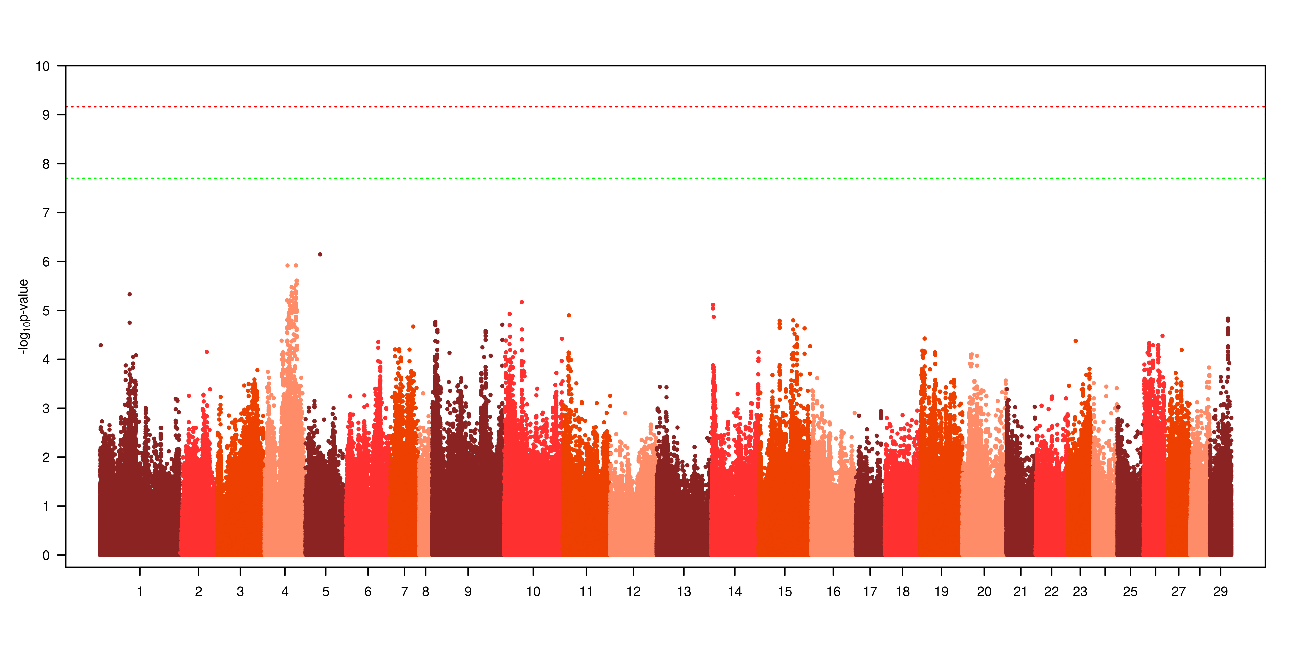

**Pop3_LC1**

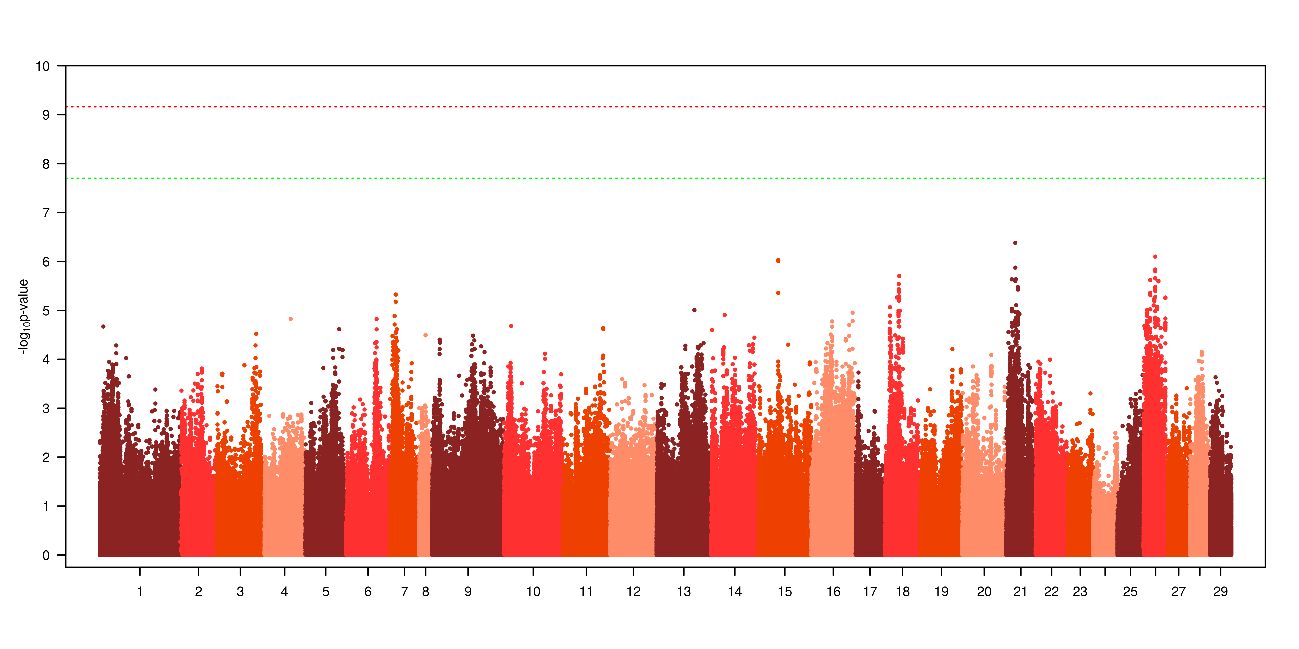

**Pop3_LC2**

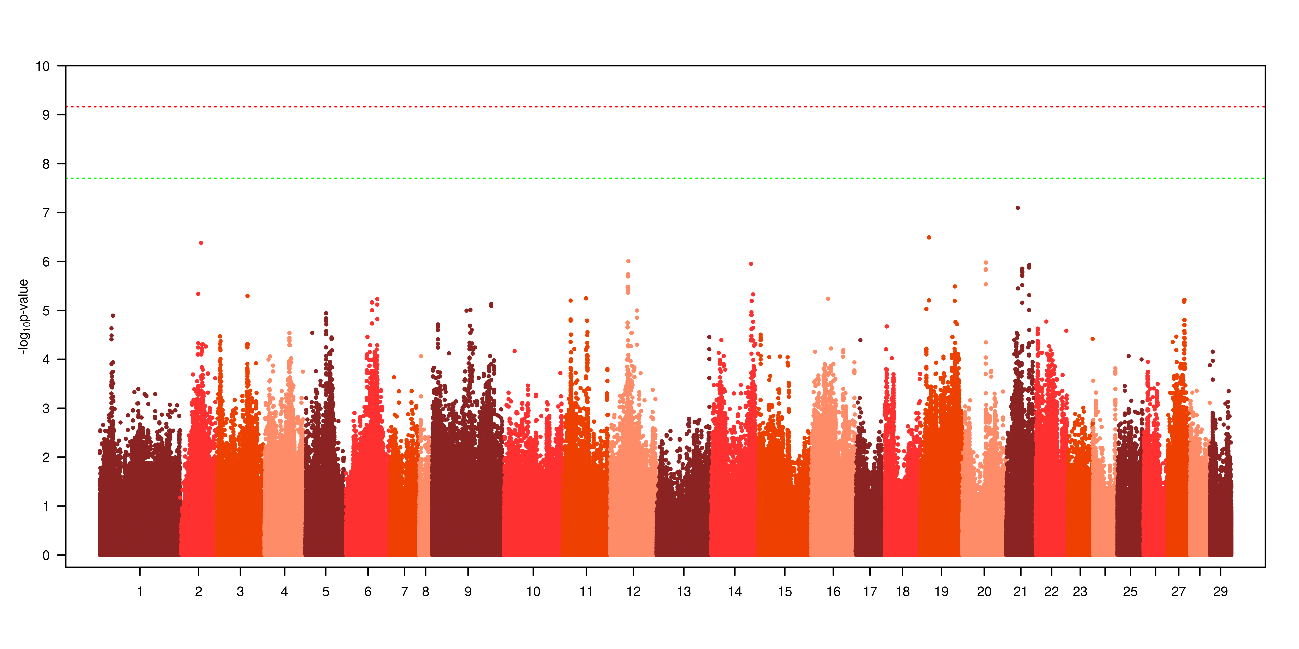

**Pop4_LC1_tk1**

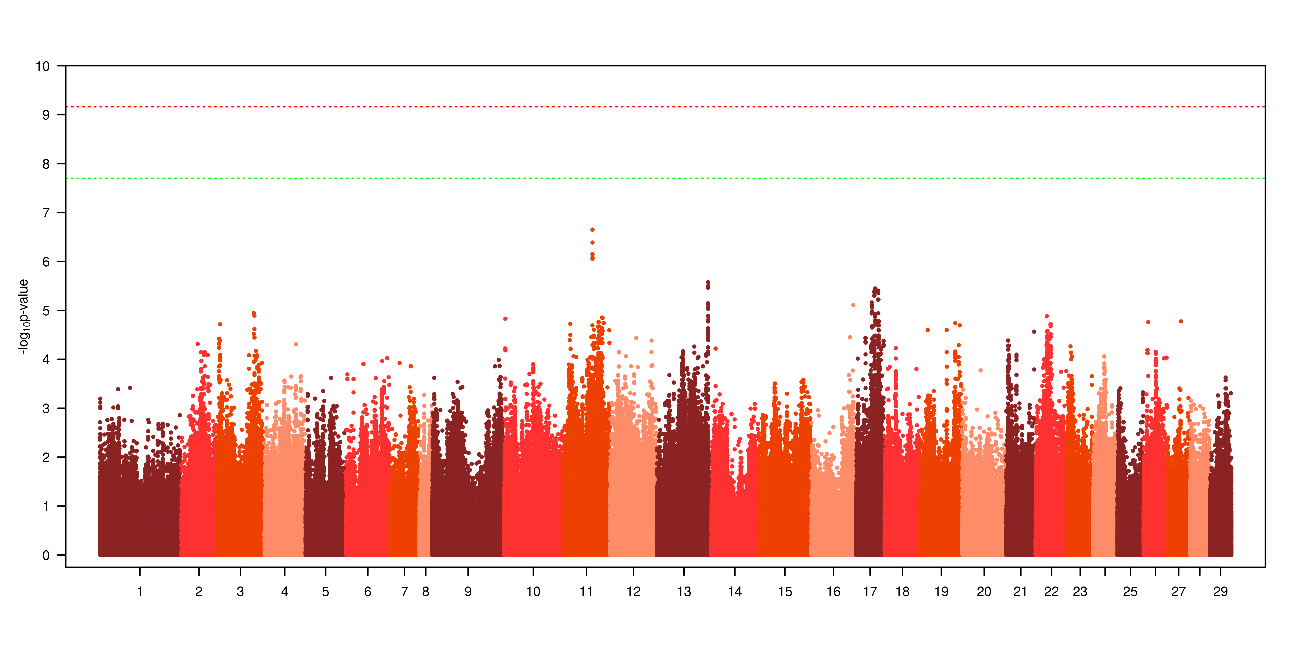

**Pop4_LC1_tk2**

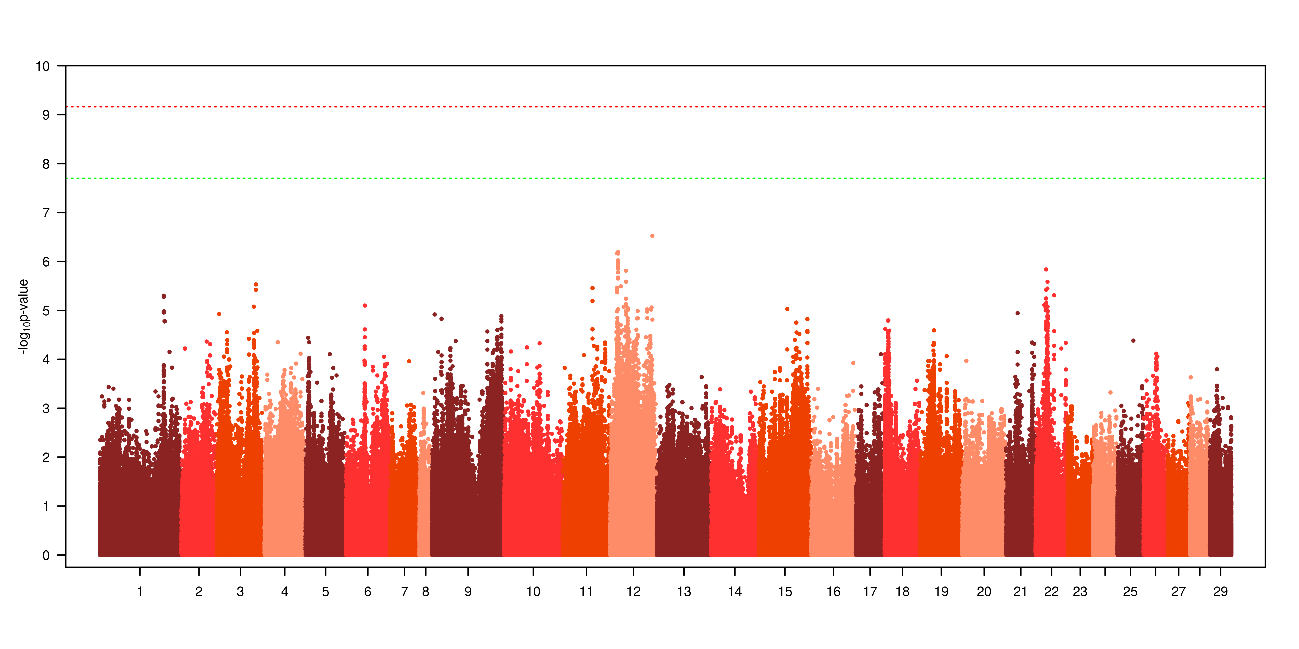

**Pop4_LC2_tk1**

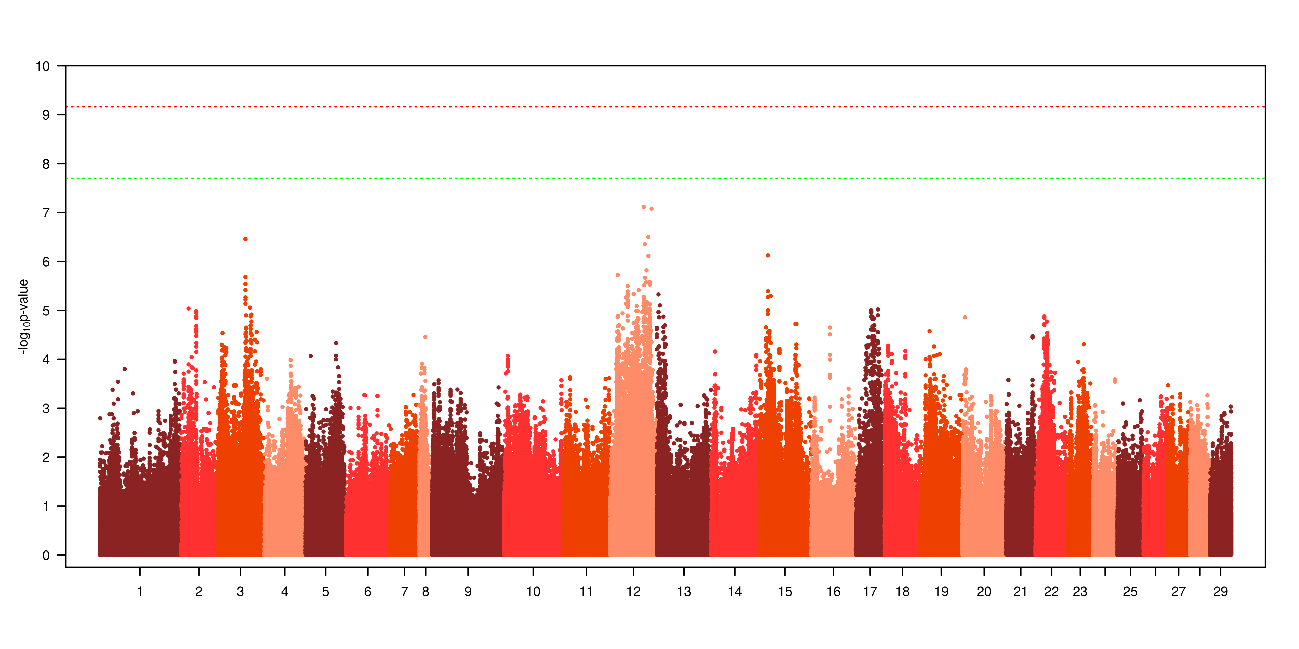

**Pop4_LC2_tk2**

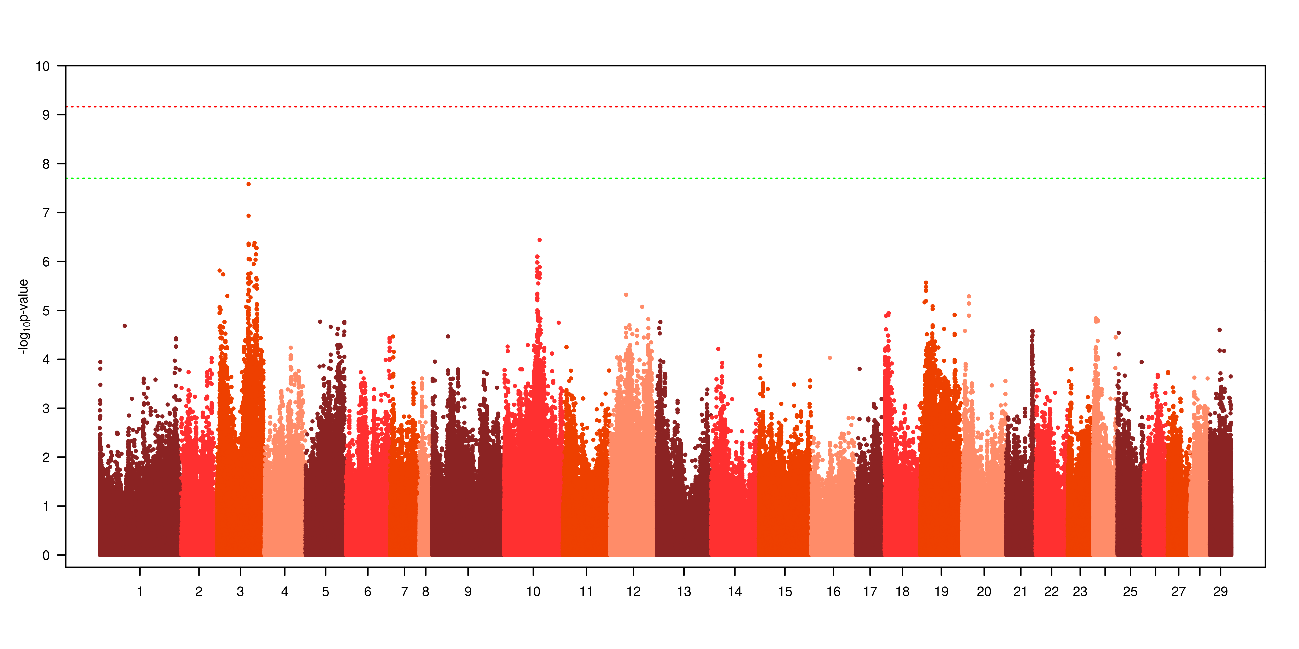

**Pop5_LC1_t1**

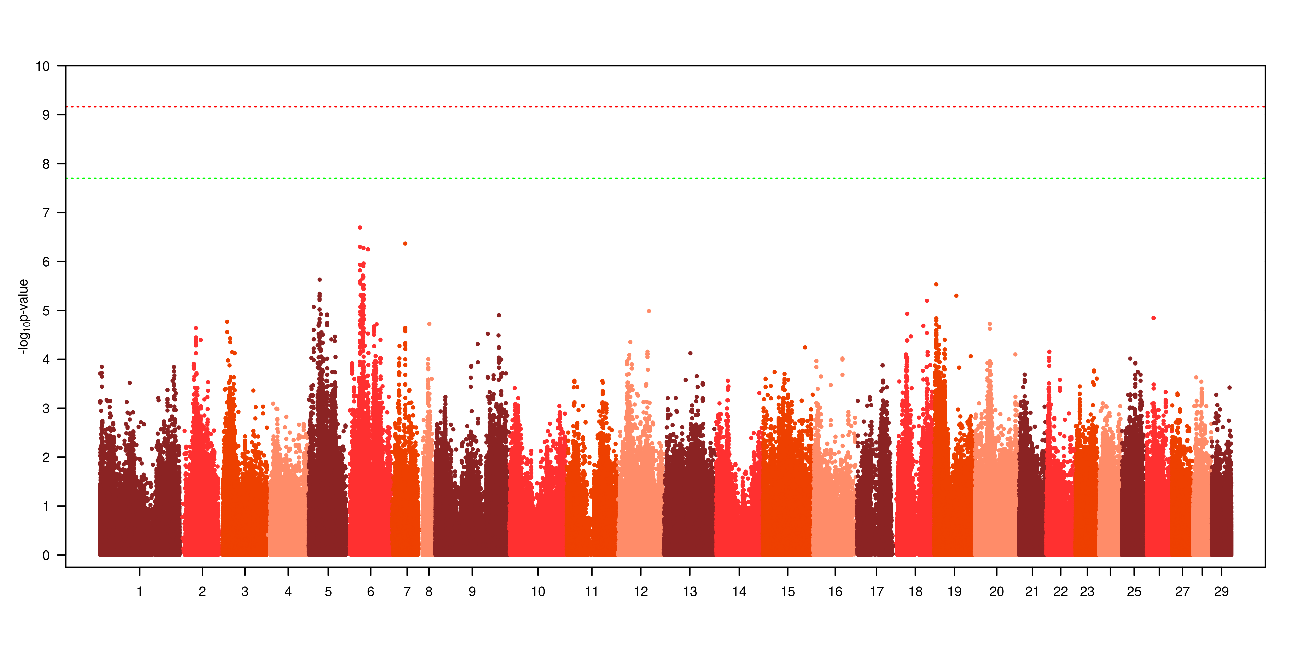

**Pop5_LC1_t2**

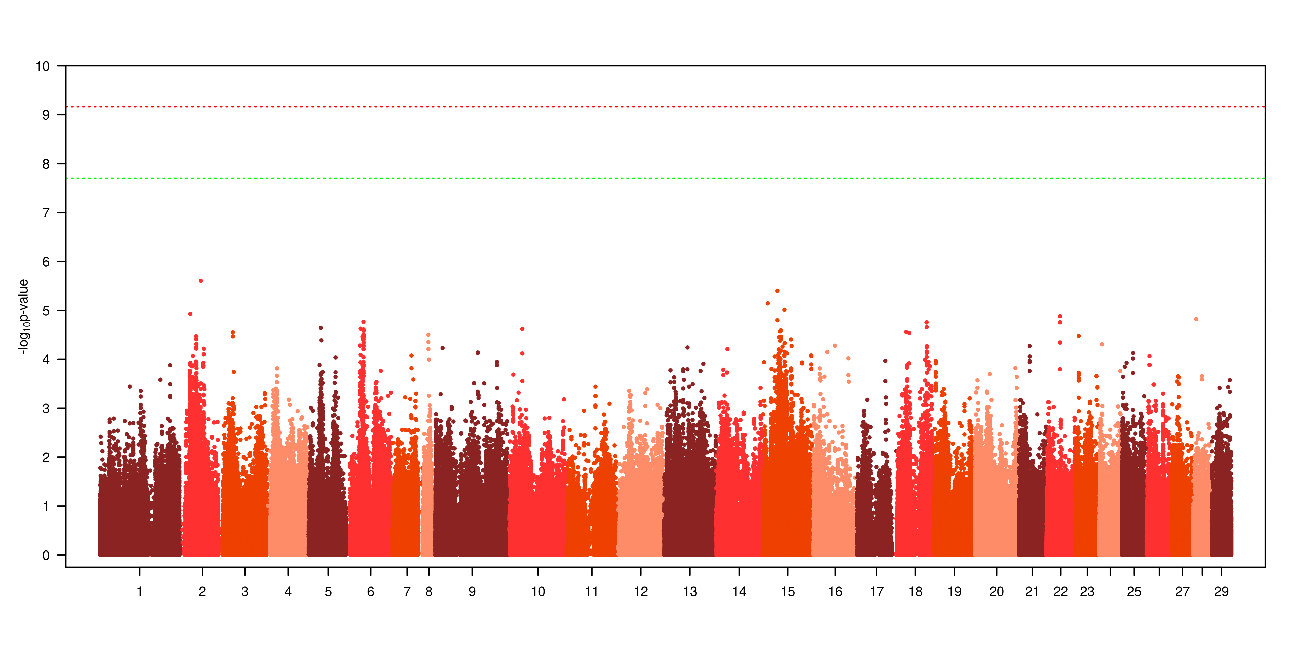

**Pop5_LC1_t3**

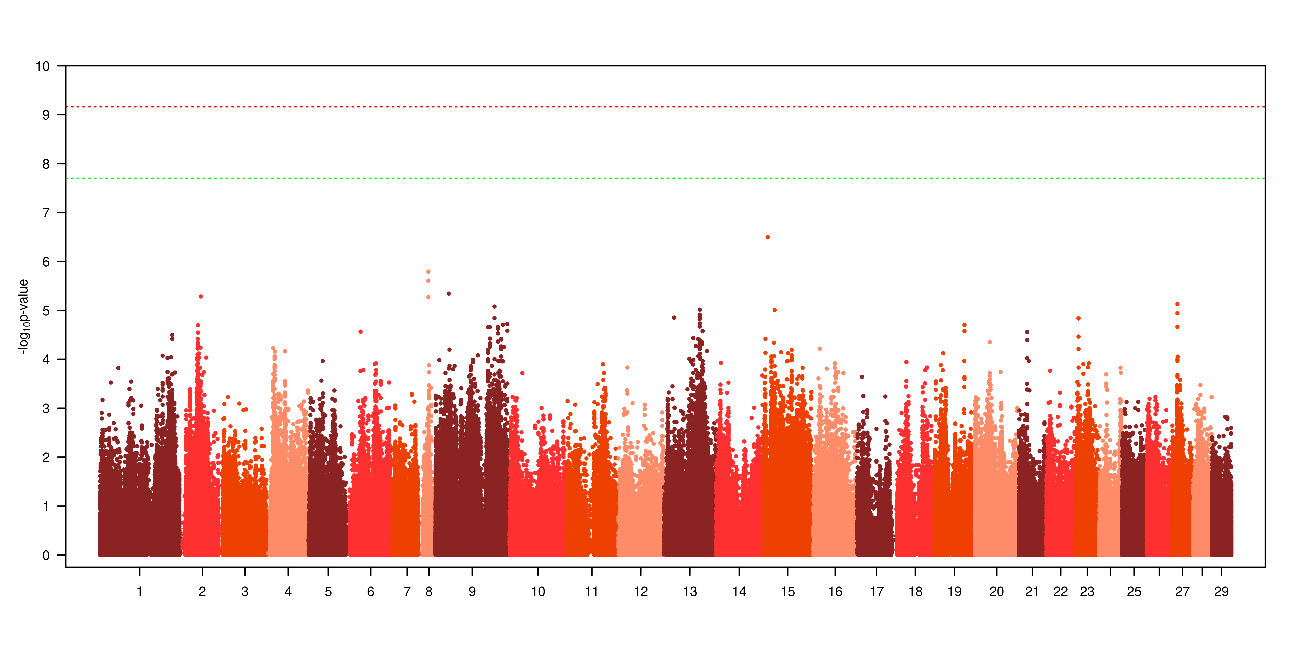

**Pop5_LC2_t1**

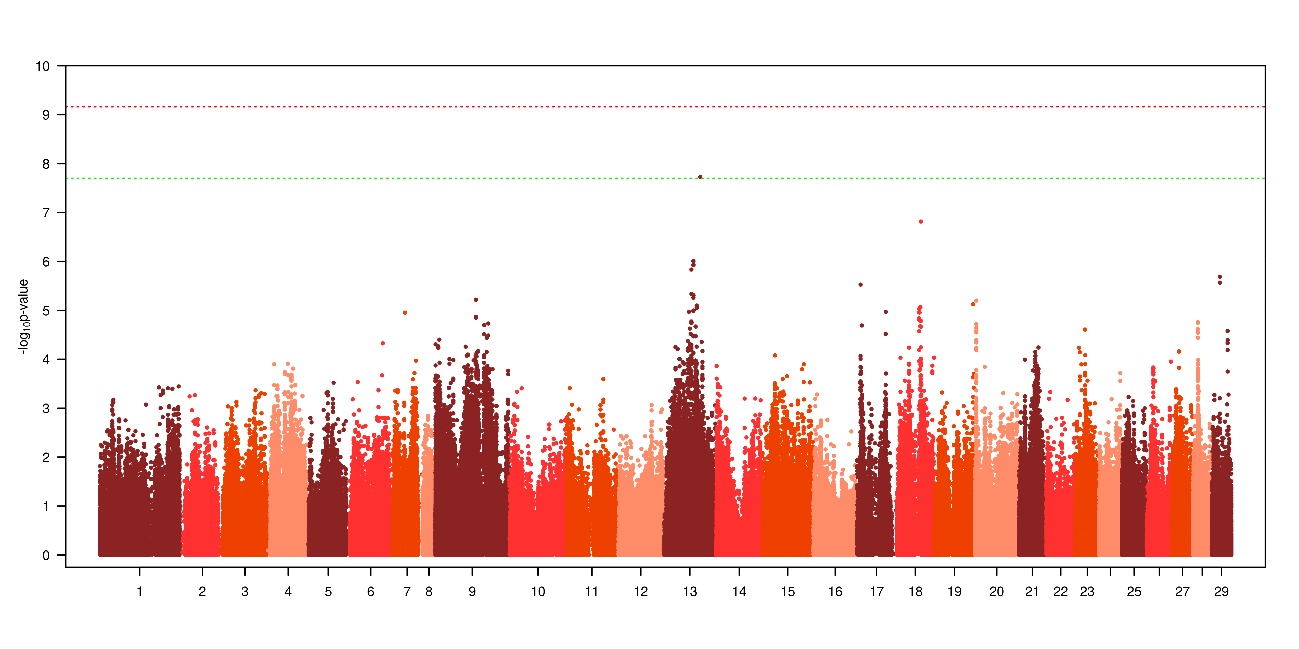

**Pop5_LC2_t2**

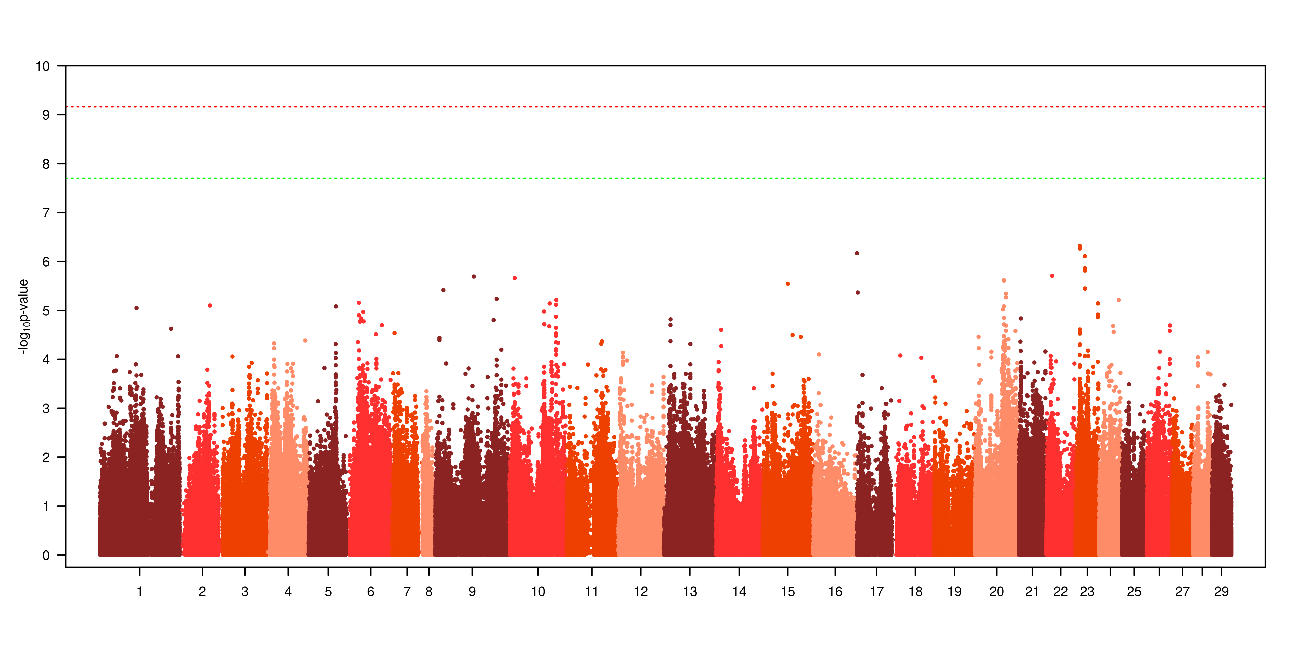

**Pop5_LC2_t3**

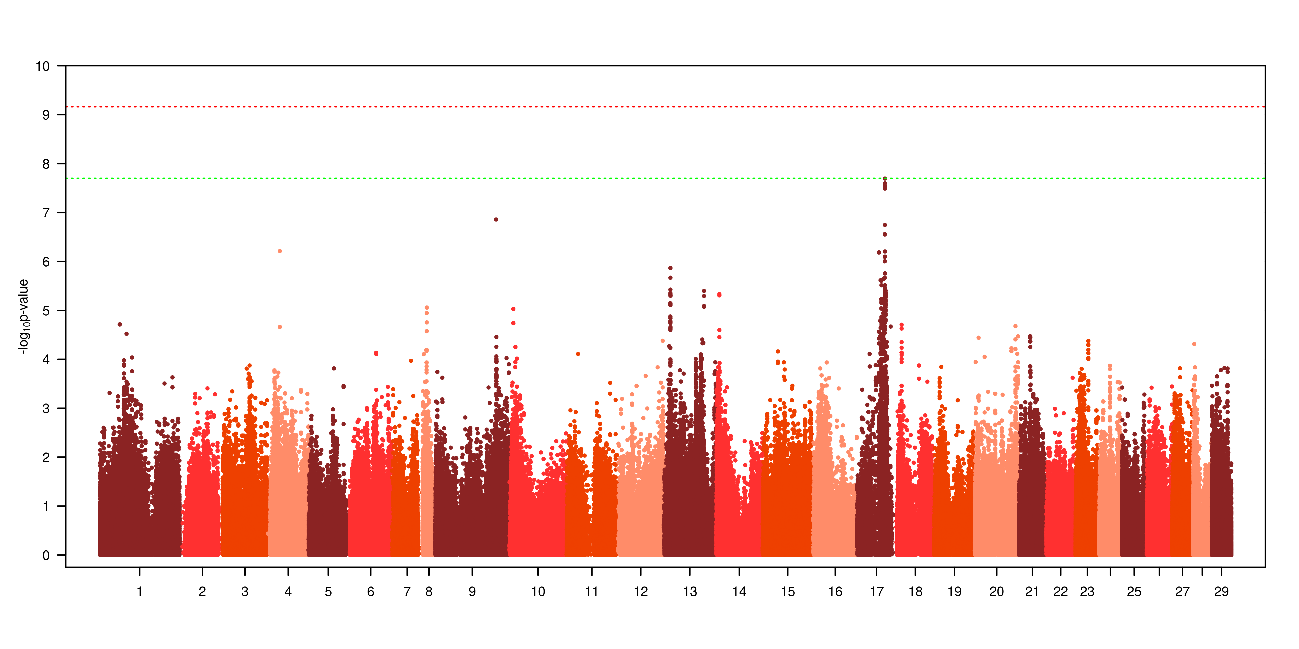
