## Supplemental 3 for "Prioritized imputed sequence variants from multi-population GWAS improve prediction accuracy for sea lice count in Atlantic salmon (*Salmo salar*)"

**Supplementary 3.** Manhattan plot of meta-analysis of GWAS for SNP prioritization for Pop4 (including only Pop1, 2, 3 and 5) and Pop5 (including only Pop1, 2, 3 and 4).

**Pop4**

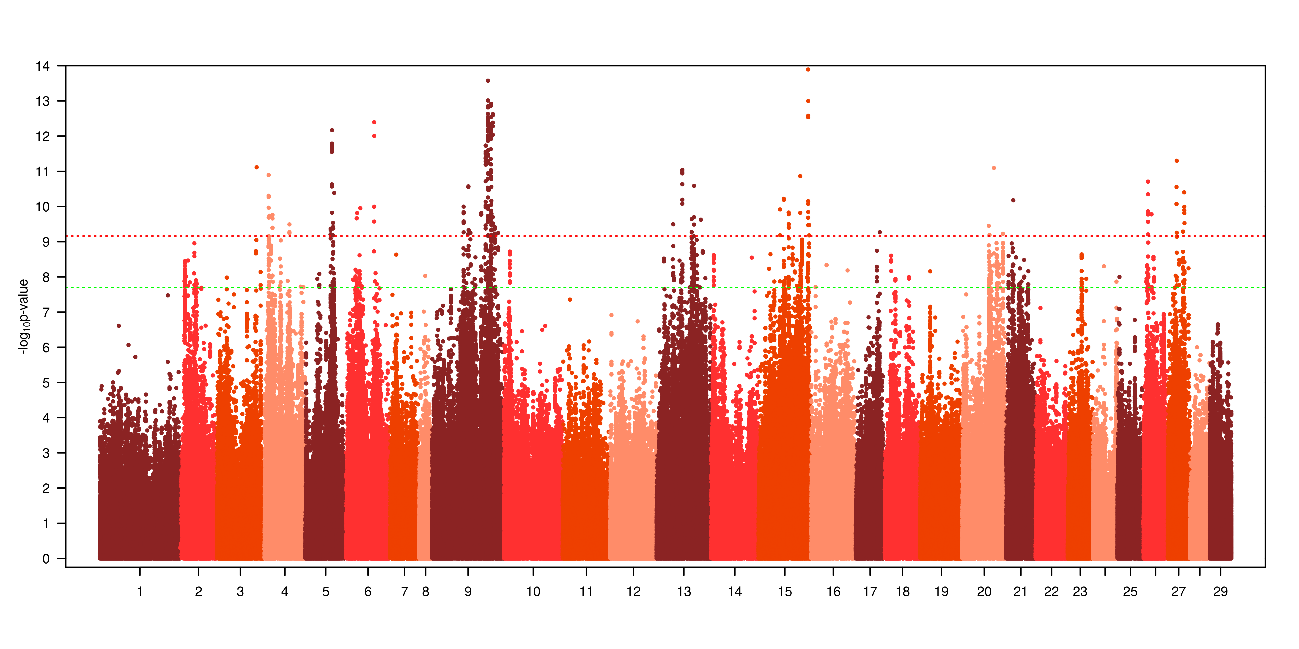

**Pop5**

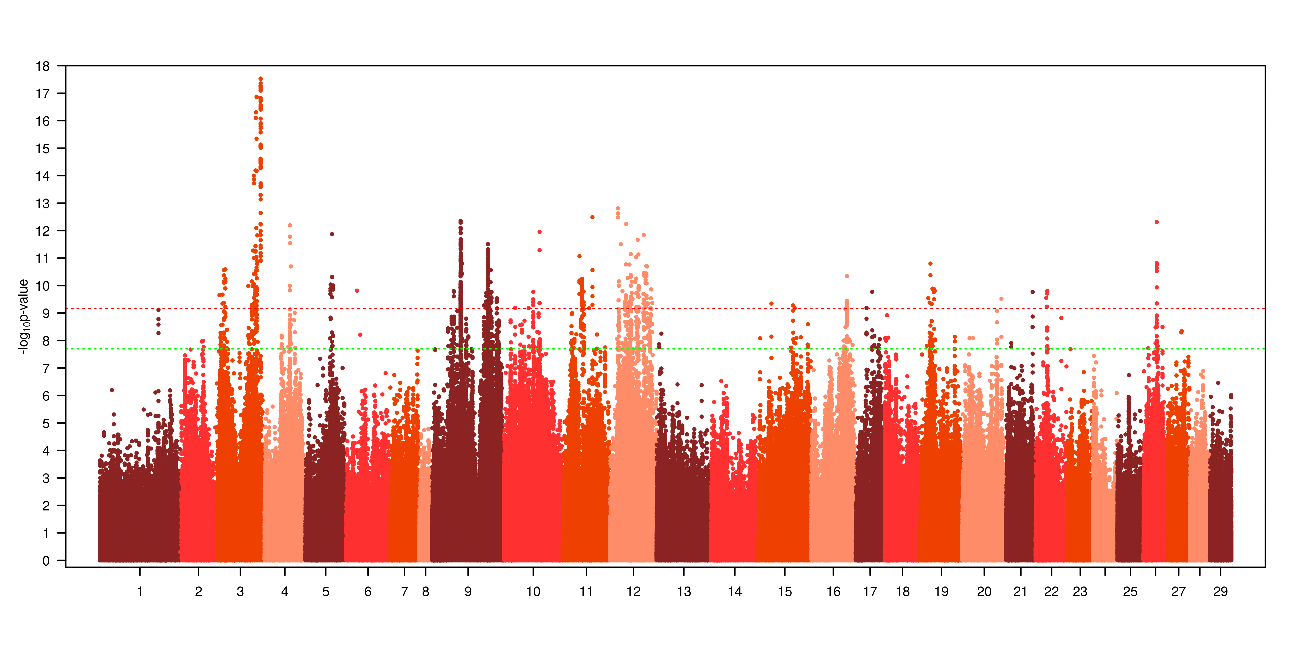
